# SPDEF promotes the classical subtype of pancreatic ductal adenocarcinoma

**DOI:** 10.1101/2022.03.18.484951

**Authors:** Claudia Tonelli, Georgi N. Yordanov, Yuan Hao, Astrid Deschênes, Olaf Klingbeil, Hsiu-Chi Ting, Erin Brosnan, Abishek Doshi, Youngkyu Park, Christopher R. Vakoc, Jonathan Preall, David A. Tuveson

## Abstract

Pancreatic ductal adenocarcinoma (PDA) samples reveal extensive cellular heterogeneity. Using single-cell RNA sequencing, we uncover multiple tumor cell populations distinguished by their differentiation state and associated with different stages of tumor progression in a mouse model of PDA. We identify Spdef as a factor required for tumorigenesis in pancreatic cancer cells of epithelial and mucinous nature. By comparative analysis of cell differentiation states in mice and humans, we find that the Spdef program is highly expressed by human PDAs of the classical subtype. Mouse and human PDA cells expressing elevated levels of Spdef are dependent upon this transcription factor for tumor progression *in vivo*. The tumor-promoting function of Spdef is recapitulated by two Spdef target genes that regulate protein folding and endoplasmic reticulum activity, Agr2 and *Ern2*/Ire1β. These findings offer insights into the factors controlling differentiation states in PDA and identify new vulnerabilities in the most common subtype of pancreatic cancer.

## INTRODUCTION

Pancreatic ductal adenocarcinoma (PDA) is a lethal cancer with a 5 year survival rate of only 11% (1). The main factors that contribute to this poor prognosis are its aggressive and metastatic nature, late diagnosis as a consequence of lack of early detection strategies and absence of specific symptoms at early stages, and a poor response to therapy (2,3).

The genetic drivers of PDA are well-described: oncogenic KRAS mutations in the exocrine pancreas serve as the initiating event and promote the formation of precursor lesions called pancreatic intraepithelial neoplasia, or PanINs. Subsequent inactivating mutations of tumor suppressor genes (e.g., *TP53, CDKN2A, SMAD4*) drive tumor progression, a finding supported by genetically engineered mouse models (GEMMs) of PDA and genetic analysis of human tumor samples (4-6). While the genetic progression of PDA is well-characterized, the underlying molecular mechanisms are less understood.

Transcriptomic studies have revealed that PDAs can be clustered into two major subtypes, termed classical and basal-like, with characteristic histopathologic features and different prognoses (7-11). PDAs of the classical subtype are more common and include well-differentiated tumors with a favorable prognosis, whereas PDAs of the basal-like subtype are poorly differentiated tumors with a worse clinical outcome (9,12). Classical PDAs present glandular structures with high secretory activity, whereas basal-like PDAs show a prevalence of undifferentiated tumor component with a dominance of mesenchymal/squamous cells. In accordance with the tumor histology, basal-like PDAs are characterized by the expression of genes involved in epithelial–to-mesenchymal transition (EMT), response to hypoxia and activation of the transcription factors MYC and ΔNp63; whereas classical PDAs are enriched for the expression of endoderm specification genes, such as HNF1A, HNF4A and GATA6 (8,9,11). Subsequent studies have demonstrated that basal-like and classical phenotypes coexist within a single tumor and that the acquisition of basal-like features results from a combination of genomic aberrations, microenvironmental changes and epigenetic factors (13,14). More recently, PDA cells that are intermediate for the expression of basal-like and classical genes have been identified (15). These cells may represent the transition state between classical and basal-like differentiation.

Identification of other factors determining these cellular phenotypes is important for understanding the mechanisms of pancreatic cancer initiation and progression and may lead to the discovery of subtype-specific dependencies. Several lineage specifiers controlling differentiation in PDA have been identified. These include, among others, KLF5, ELF3 and FOXA1 for the classical subtype and TP63, ZEB1, ZBED2 and GLI2 for the basal-like subtype (16-20).

In this study, we reveal various pancreatic cancer cell subpopulations characterized by different differentiation states based on their gene expression profiles. We find that pancreatic cancer cells of epithelial and mucinous nature express SAM pointed domain containing ETS transcription factor (Spdef or prostate-derived ETS factor Pdef). SPDEF is the sole member of the class IV of ETS family of transcription factors because of its unique binding preference for a GGAT rather than a GGAA core (21,22). The expression of Spdef is largely restricted to epithelial tissues including the lung, stomach and intestine, where it is essential for the maturation of Goblet and Paneth cells (23-27). In secretory cells, Spdef regulates a network of genes associated with mucus production, protein folding and glycosylation (23,28-31).

In epithelial tissues such as the breast, prostate, lung and colon, Spdef plays a role in tumorigenesis. In prostate cancer, Spdef reduced metastasis formation by inhibiting cell migration and invasion (32-35). Similarly, in colorectal cancer, Spdef hampered tumor progression and loss of Spdef enhanced tumor formation and led to local invasion (36,37). Re-expression of SPDEF in colon cancer cells inhibited growth and migration (38). In breast cancer, Spdef exhibited both tumor-suppressive and tumor-promoting functions. SPDEF expression was lost in invasive breast cancer cell lines and its re-expression inhibited the proliferation and the migration of these cells (39,40). In contrast, ectopic expression of SPDEF induced invasion and cell motility in mammary epithelial cells and enhanced their tumorigenic growth in immunocompromised mice (41,42). In addition, SPDEF was shown to be overexpressed in breast cancer compared to normal tissue and act as a survival factor in ER-positive breast cancer (43-46). Finally, in lung cancer, Spdef cooperated with mutant Kras to induce malignant mucinous lung tumors (47).

In this study, we sought to investigate the role of SPDEF in pancreatic cancer progression. We hypothesize that SPDEF may be selectively involved in the epithelial cell fate specification and survival of well-differentiated and highly secretory PDAs of the classical subtype.

## RESULTS

### Single-cell RNA sequencing reveals the transcriptional dynamics of pancreatic cancer progression

To investigate pancreatic cancer progression, we utilized the *Kras*^*LSLG12D/+*^; *Trp53*^*LSLR172H/+*^; *Pdx1-Cre* (KPC) GEMM in which Kras^G12D^ and p53^R172H^ were expressed in the pancreas by virtue of a Pdx1-Cre transgene (48,49). KPC tumors are characterized by intra-tumor histological heterogeneity: areas of acinar-to-ductal metaplasia (ADM), low-grade and high-grade PanINs are intermixed with invasive cancer cells (48,49). To unravel the transcriptional dynamics underlying tumor progression that result in intra-tumor heterogeneity, we performed single-cell RNA sequencing (scRNA-seq) on tumor cells isolated by negative selection from 8 KPC tumors (Figure S1A). Clustering of batch-adjusted, combined data of 23,991 KPC tumor cells identified 6 distinct cell populations, with similar contribution of cells from each tumor (Figure S1B, C).

Three of these cell populations were clearly identifiable: “epithelial^high^” cancer cells in cluster 1 represented 22% of all tumor cells and expressed high levels of the epithelial markers *Krt18, Krt8, Cdh1* and low levels of the mesenchymal marker *Vim*; mesenchymal cancer cells in cluster 3 expressed genes associated with a mesenchymal differentiation (*Twist1, Yap1, Vim*) and represented 12% of all tumor cells; finally, proliferating cancer cells in cluster 5 expressed genes encoding for proteins involved in cell cycle and mitosis (*Cdk1, Ube2c, Cenpa*) and represented 5% of all tumor cells (Figure 1A and S1D). Cluster 0, 2 and 4 were not readily identifiable.

**Figure 1.**
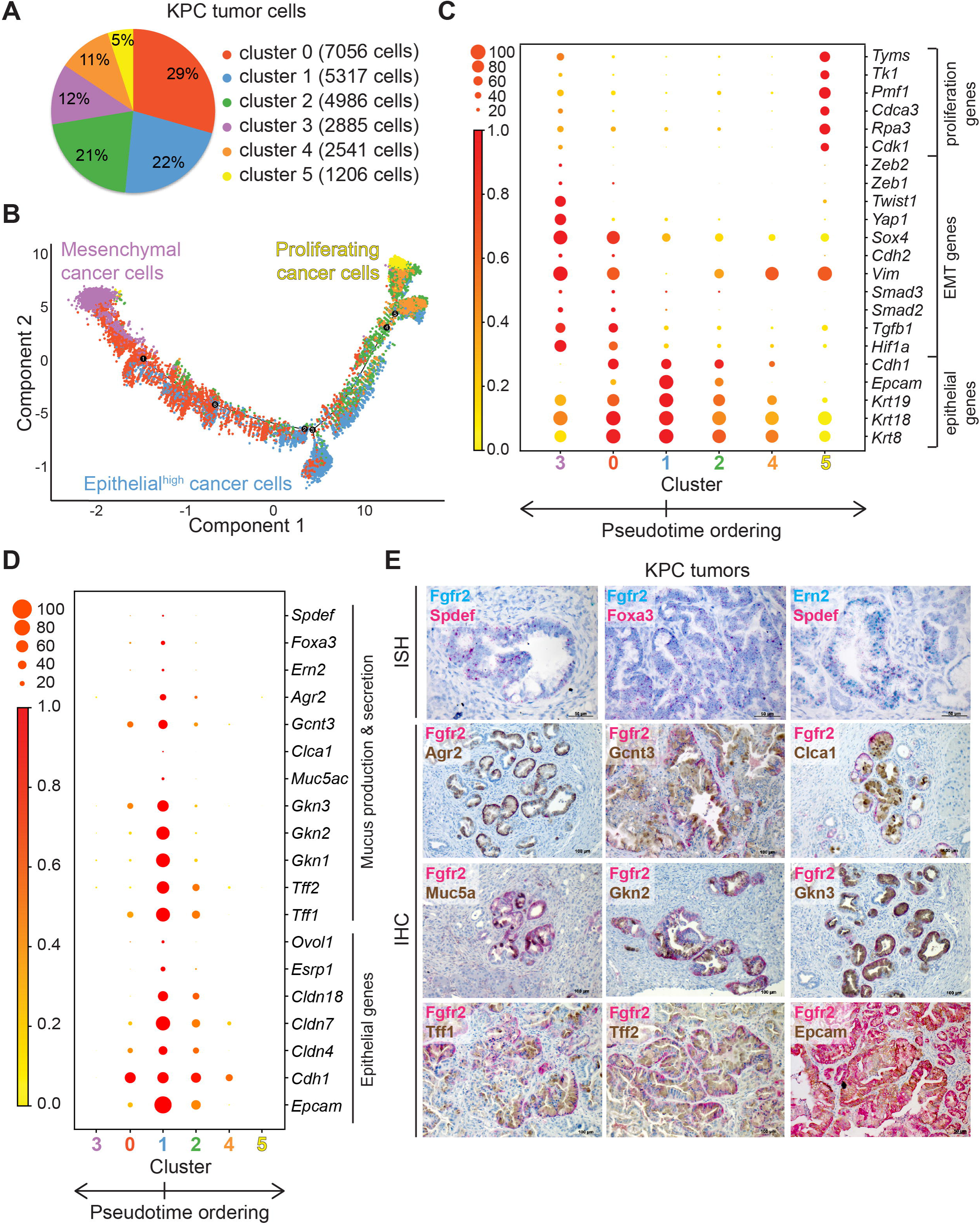
Single-cell RNA sequencing reveals the transcriptional dynamics of pancreatic cancer progression. **A**. Pie chart showing the distribution of cells in each cluster. **B**. Pseudotime ordering of cancer cells, coloring by cluster. **C, D**. Dot plot of the expression of the indicated genes in the different clusters. The size of each dot represents the percentage of cells within a given cluster that expresses the gene; the intensity of color indicates the level of mean expression. The order of the clusters matches the order of the clusters in the pseudotime trajectory. **E**. Representative RNA ISH of *Fgfr2* (blue staining), *Spdef* (red staining), *Foxa3* (red staining), *Ern2* (blue staining) and immunohistochemical staining for Fgfr2 (red staining) and Agr2, Gcnt3, Clca1, Muc5ac, Gkn2, Gkn3, Tff1, Tff2, Epcam (brown staining) in KPC tumor sections.

To reveal the evolutionary relationships among these populations, we ordered cells in pseudotime based on their transcriptional similarity (50). This unsupervised analysis ordered the cells on a V-shaped timeline and placed the epithelial^high^ cancer cells at the bottom of the V and the mesenchymal and proliferating cancer cells at the opposite ends of the trajectory (Figure 1B). The epithelial^high^ cancer cells expressed *Fgfr2, Mmp7* and *Lcn2*, which we previously identified as markers of pre-invasive cancer cells in KPC tumors (Figure S1E) (51). Thus, we interpreted this trajectory as paths of disease progression in which the epithelial^high^ cancer cells acquired either a mesenchymal or proliferative phenotype. During malignant differentiation, cells transitioned through different states represented by clusters 0, 2 and 4. The mesenchymal or proliferative phenotypes were likely not end states, however subsequent evolutionary changes were not captured in this analysis.

We next explored how expression states changed during malignant differentiation. Epithelial^high^ cancer cells in cluster 1 expressed high levels of the epithelial markers *Cdh1, Epcam, Krt19, Krt18* and *Krt8* and their expression was progressively lost or decreased along the pseudotime branches (Figure 1C). Cells on the left branch of the pseudotime trajectory progressively acquired the expression of the master-regulator of hypoxic signaling *Hif1a* and the genes in the TGFβ pathway *Tgfb1, Smad2* and *Smad3*. Both Hif1α and TGFβ are known regulators of EMT (52-57). Indeed, increased activation of the Hif1α and TGFβ pathway corresponded with increased expression of genes associated with EMT, such as *Vim, Cdh2, Sox4, Yap1, Twist1, Zeb1* and *Zeb2* (58-60). The highest expression of these genes was achieved in the mesenchymal cancer cells in cluster 3. In addition to the induction of EMT, TGFβ is known to induce switching of FGF receptors in epithelial cells (61). Indeed, epithelial^high^ cancer cells progressively lost the expression of the epithelial receptor *Fgfr2* and acquired the expression of the mesenchymal receptor *Fgfr1* as they transitioned from an epithelial to a more aggressive mesenchymal phenotype (Figure S1F) (62). Cells on the right branch of the pseudotime trajectory also lost or decreased the expression of epithelial markers and acquired the expression of the mesenchymal marker *Vim*. However, different from the cells on the left branch of the trajectory, cells on the right branch of the trajectory did not activate the Hif1α or TGFβ pathway and exhibited a partial EMT phenotype. Among these partial EMT cancer cells, the ones in cluster 5 were actively replicating and expressed several genes involved in cell cycle, mitosis and nucleotide biosynthesis, such as *Cdk1, Rpa3, Cdca3, Pmf1, Tk1* and *Tyms*.

In summary, epithelial^high^ cancer cells progressed to more aggressive phenotypes by losing epithelial features, activating a partial or complete EMT program and finally acquiring mesenchymal or proliferative phenotypes both of which have been described in human PDAs (63,64). Our analysis uncovered the expression states underlying malignant differentiation in pancreatic cancer, with implications for the human disease.

### Epithelial^high^ pancreatic cancer cells express genes associated with mucus production and secretion

To further investigate the biology of KPC epithelial^high^ cancer cells, we determined the marker genes for cluster 1 (Table S1). In addition to strong expression of epithelial genes, we found that epithelial^high^ cancer cells up-regulated genes associated with mucus production and secretion, including the transcription factors *Spdef* and *Foxa3* (Figure 1D).

In secretory cells, Spdef activates Foxa3 and they regulate a network of genes associated with mucus production, protein folding and glycosylation, including the ER stress sensor *Ern2*/Ire1β, the disulfide isomerase Agr2, the glycosylation enzyme Gcnt3, the chloride channel involved in mucus production Clca1 and the mucin Muc5ac, among others (23,28-31).

Here, we found that this secretory cell gene signature was induced in KPC epithelial^high^ cancer cells. In addition, we observed the expression of gastric tissue genes, such as the gastrokines (Gkns) and the trefoil factors (Tffs) (65,66). The activation of these signatures was previously observed during pulmonary to gastric transdifferentiation of neoplastic lung cells following loss of the pulmonary lineage specifier Nkx2.1 and during the oncogenic Kras-induced metaplastic development of pancreatic cancer precursors (67-71).

We analyzed the expression of *Spdef* and *Foxa3*, the secretory cell genes *Ern2*, Agr2, Gcnt3, Clca1, Muc5ac, the gastric genes Gkn2, Gkn3, Tff1, Tff2 and the epithelial genes Epcam and Fgfr2 by immunohistochemistry (IHC) or RNA in situ hybridization (RNA ISH) in KPC tumor sections (Figure 1E). We found that these genes were expressed in glandular lesions comprised of columnar, mucinous neoplastic cells of epithelial nature. As these epithelial^high^ cancer cells progressed in the disease, they lost the expression of these genes, became more disorganized, stopped producing mucus and acquired invasive features, in accordance with the tumor progression model inferred from the analysis of the scRNA-seq data.

### Spdef and its target genes *Ern2*/Ire1β and Agr2 support murine pancreatic tumor growth

The role of Spdef and Foxa3 in tumorigenesis is unclear, exhibiting both tumor-suppressive and tumor-promoting functions. To investigate their role in pancreatic cancer progression, we inactivated *Foxa3* and *Spdef* expression in mouse epithelial tumor organoids (‘mT’) using small guide RNA (sgRNA) pairs designed to delete the transcription start site (TSS) and isolated single cell-derived clones. *Foxa3* and *Spdef* loss was assessed at the mRNA level, as we could not identify reliable antibodies (Figure S2A-D). The results on the effect of *Foxa3* deletion on the growth of mT69a organoids *in vivo* did not demonstrate a dependency upon tumor progression: one *Foxa3* knock-out clone formed tumors as efficiently as the control clone, while the other *Foxa3* knock-out clone progressed faster than the control clone (Figure 2A; S2E, F). On the contrary, *Spdef* inactivation in several clones of mT69a severely impaired tumor growth *in vivo* following orthotopic transplantation, and extended the survival of these orthotopically grafted organoid models (Figure 2B, C; S2G, H). The tumor-promoting role of Spdef in pancreatic cancer progression was also confirmed in the tumor organoid line mT6 (Figure 2D; S2I, J). Notably, *Spdef* inactivation did not have a clear effect on tumor organoid proliferation *in vitro* and changes in duplication rate were minor and more likely due to clonality (Figure S2K).

**Figure 2.**
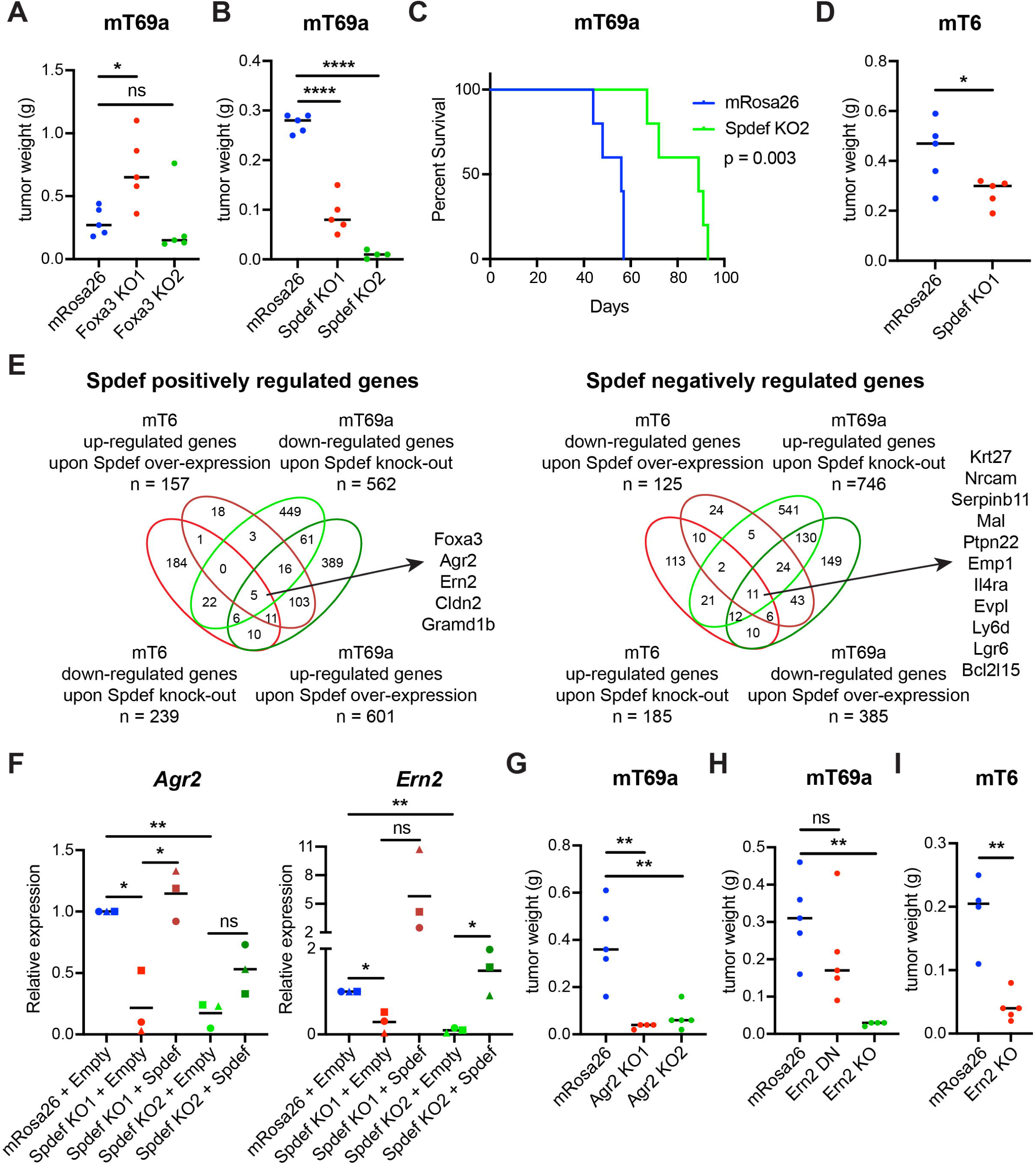
Spdef and its target genes *Ern2*/Ire1β and Agr2 support murine pancreatic tumor growth. **A, B, D, G, H, I**. Quantification of weight of tumors derived from mT69a (**A, B, G, H**) and mT6 (**D, I**) orthotopically grafted organoid (OGO) models of *Foxa3* knock-out (KO) (**A**), *Spdef* KO (**B, D**), *Agr2* KO (**G**), *Ern2* KO or partial inactivation DN (**H, I**) and mRosa26 clones in nu/nu mice. Results show mean of biological replicates. Unpaired Student’s t test. **C**. Kaplan–Meier survival curve of percent survival for mT69a OGO models of *Spdef* KO (n=5) and mRosa26 (n=5) clones in nu/nu mice. Log-rank (Mantel–Cox) test. **E**. Venn diagram of the differentially expressed genes identified by RNA-seq following *Spdef* KO and restoration by cDNA expression in mT69a and mT6 organoids (up-regulated genes: q-value < 0.05, log2 of fold change > 0; down-regulated genes: qvalue < 0.05, log2 of fold change < 0). **F**. *Agr2* and *Ern2* expression as determined by RT-qPCR in mT69a (triangle), mT6 (circle) and mT23 (square) organoids following *Spdef* KO and re-expression. Paired Student’s t test.

To determine the mechanism underlying the tumor growth defect observed upon Spdef loss in mouse epithelial tumor organoids following orthotopic transplantation, we performed RNA sequencing (RNA-seq) of mT6 and mT69a organoid clones with or without knock-out of *Spdef* and restoration by cDNA expression (Figure 2E; Table S2). We investigated the Spdef-regulated genes whose differential expression upon *Spdef* deletion was reverted by Spdef re-expression. Of these genes, 5 were down-regulated upon *Spdef* knock-out and up-regulated upon its re-expression, while 11 were up-regulated upon *Spdef* knock-out and down-regulated upon its re-expression in both mT6 and mT69a organoids. The positively regulated genes were the transcription factor *Foxa3*, the ER-resident disulfide isomerase *Agr2*, the ER stress sensor *Ern2*/Ire1β, the tight junction protein *Cldn2* and the cholesterol transporter *Gramd1b*.

Spdef was highly expressed in mucinous epithelial^high^ cancer cells in KPC tumors (see Figure 1E). Mucus-secreting cells rely on ER activity to achieve proper folding of secreted proteins and prevent ER stress (72,73). As both Agr2 and *Ern2*/Ire1β are localized in the ER and play a role in the maintenance of ER homeostasis (74-76), we next investigated whether deletion of *Agr2* and *Ern2* would mimic the effect of deletion of *Spdef*. We confirmed by RT-qPCR in mT6, mT23 and mT69a organoids that *Agr2* and *Ern2* were regulated by Spdef as previously reported (Figure 2F) (23,28). *Agr2* and *Ern2* inactivation was achieved with sgRNA pairs designed to delete the TSS followed by isolation of single cell-derived clones. *Agr2* and *Ern2* loss was assessed at the mRNA level and by confirming the deletion of the TSS at the genomic DNA level (Figure S3A-C). We found that deletion of *Agr2* in two different mT69a clones reduced tumor growth *in vivo* following orthotopic transplantation (Figure 2G; S3D, E). Complete deletion of *Ern2* in mT69a and mT6 strongly impaired tumor growth *in vivo* (Figure 2H, I; S3F-I). Of note, partial inactivation of *Ern2* in mT69a had an intermediate effect. Thus, similar to Spdef, *Ern2*/Ire1β and Agr2 promoted the progression *in vivo* of epithelial pancreatic cancer cells.

The tumors formed following mT organoid implantation were highly cellular, poorly differentiated and did not produce mucus, independently of whether they were derived from mT organoids expressing or not *Spdef, Agr2* or *Ern2* (Figure S2H, J and S3E, G, I). We analyzed *Spdef* and *Ern2* expression by RNA ISH and Agr2 expression by IHC and found that most of the malignant cells in control tumors did not express *Spdef, Ern2* or Agr2 except for few rare lesions (Figure S3J). We concluded that Spdef, *Ern2*/Ire1β and Agr2 were required in the early events of tumorigenesis before tumor cells lost their epithelial and secretory nature and underwent malignant differentiation into more invasive phenotypes.

### Expression of the SPDEF program is associated with classical differentiation in PDA

To identify the human PDA cells that are in the same differentiation state as the KPC epithelial^high^ cancer cells, we evaluated the expression of a humanized version of the marker genes up-regulated in scRNA-seq cluster 1 in RNA-seq data of laser-capture microdissected (LCM) epithelium from patients with PDA (77). We found higher expression of the marker genes in PDA samples of the classical subtype than of the basal-like subtype, consistent with the epithelial nature of classical tumors (Figure 3A). Of note, 8 of the marker genes of KPC epithelial^high^ cancer cells were also markers of the classical subtype, while only 2 were markers of the basal-like subtype (9).

**Figure 3.**
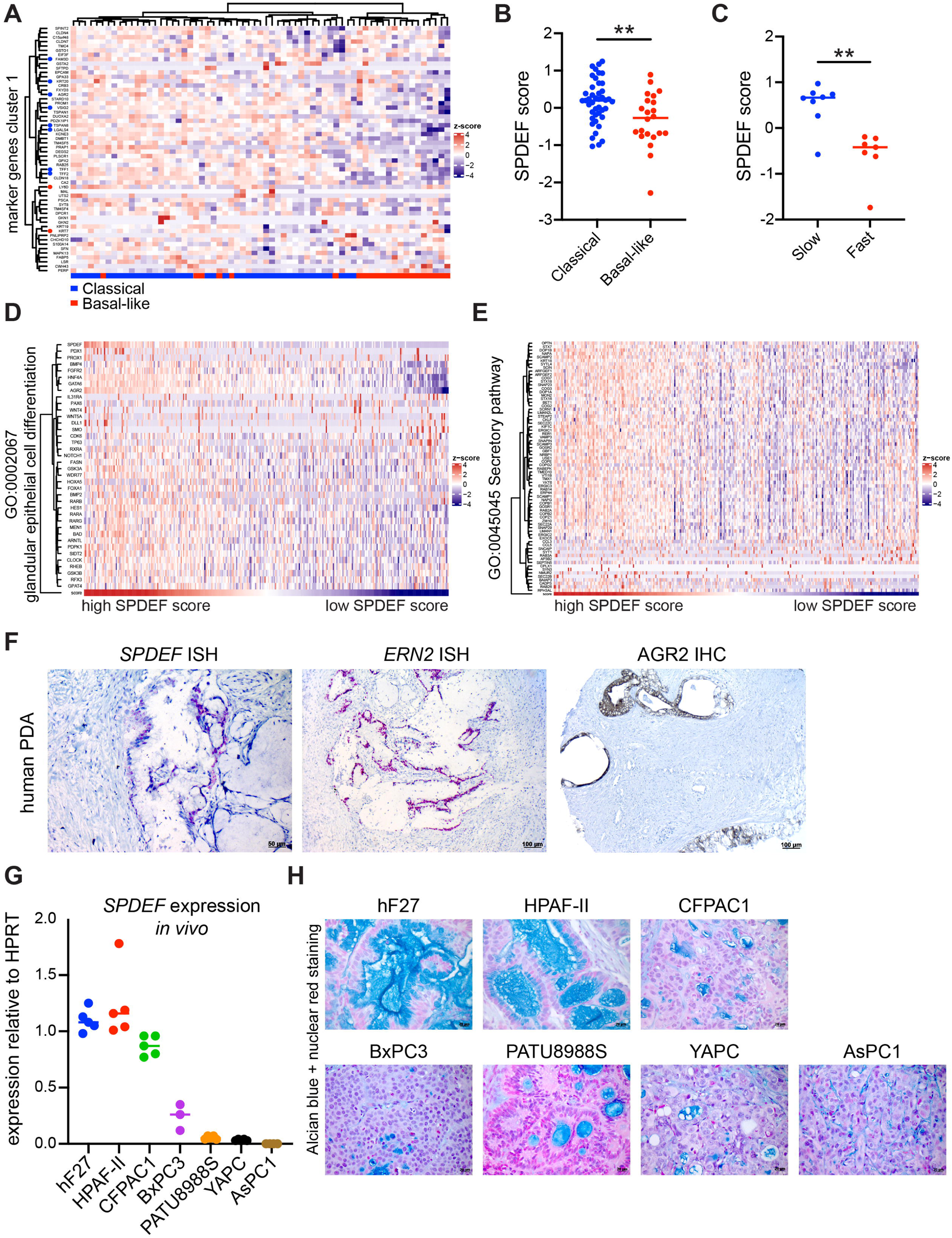
Expression of the SPDEF program is associated with classical differentiation in PDA. **A**. Heatmap showing unbiased hierarchical clustering of RNA-seq data of LCM epithelium from patients with PDA (77) for marker genes up-regulated in scRNA-seq cluster 1 (Table S1). The color scheme of the heatmap represents z-score of log_2_(normalized counts). Blue and red horizontal bar at the bottom indicates classical and basal-like subtype of PDA samples, respectively. Blue and red dots next to the gene name to the left of the heatmap indicate known classical and basal-like genes based on Moffit classification (9), respectively. **B**. SPDEF score of classical and basal-like PDA samples from (77). **C**. SPDEF score of Slow and Fast-progressing human organoid lines from (78). **D, E**. Heatmap showing unbiased hierarchical clustering of RNA-seq data from the PanCuRx Translational Research Initiative dataset of patients with locally advanced PDA (79) for genes of GO signatures ‘glandular epithelial cell differentiation’ (**D**) and ‘secretory pathway’ (**E**). Samples are ranked based on their SPDEF score. The color scheme of the heatmap represents z-score of log_2_(TPM values). **F**. Representative RNA ISH of *SPDEF* and *ERN2* and immunohistochemical staining for AGR2 in human PDA sections. **G**. RT-qPCR analysis of *SPDEF* expression in control tumors derived from transplant models of hF27, HPAF-II, CFPAC1, BxPC3, PATU8988S, YAPC, AsPC1 in NSG mice. Results show mean of biological replicates. **H**. Representative AB and nuclear red staining of control tumors derived from transplant models of hF27, HPAF-II, CFPAC1, BxPC3, PATU8988S, YAPC, AsPC1 in NSG mice.

To assess if classical PDAs expressed the SPDEF program, we calculated an SPDEF regulatory score based on the expression of *SPDEF* and its target genes *FOXA3, ERN2, AGR2* and *MUC5AC*. We established that classical tumors had a significantly higher SPDEF regulatory score than basal-like tumors (Figure 3B; S4A). To corroborate this finding, we analyzed the expression of the SPDEF program in our models of intraductally grafted slow and fast-progressing human organoid lines, which recapitulated the features of the classical and basal-like subtypes of human PDA, respectively (78). We found that slow progressors presented a significantly higher SPDEF regulatory score than fast progressors, supporting a role for SPDEF in tumors with features of the classical subtype (Figure 3C; S4B).

PDAs of the classical subtype present a dominance of well-differentiated epithelial tumor component and are characterized by the presence of glandular, mucinous lesions (12). To assess if the expression of the SPDEF regulatory program correlated with glandular epithelial differentiation and secretory activity in human PDA samples, we analyzed the RNA-seq data from the PanCuRx Translational Research Initiative dataset of patients with locally advanced PDA (79). We calculated an SPDEF regulatory score for each patient, ranked them based on their SPDEF score and plotted the expression of Gene Ontology (GO) signatures. We found that patients with high SPDEF scores presented higher expression of the GO signatures ‘glandular epithelial cell differentiation’, ‘secretory pathway’, ‘positive regulation of regulated secretory pathway’ and ‘epithelial cell differentiation’ (Figure 3D, E; S4C, D).

To evaluate if the findings from these *in silico* analyses were supported by histological investigation, we analyzed *SPDEF* and *ERN2* expression by RNA ISH and AGR2 expression by IHC in human PDA and confirmed that SPDEF and its target genes were expressed in tumor samples that presented glandular architecture and columnar mucinous epithelium (Figure 3F).

Next, we investigated *SPDEF* expression and mucus production by Alcian blue (AB) staining in several PDA cell lines defined as classical or basal-like by transcriptomic analyses (Figure S4E) (16,17,78,80). We found that *SPDEF* mRNA levels were variable in PDA cell lines and did not correlate with a secretory phenotype *in vitro* (Figure S4F, G). The only cells to stain weakly positive for mucus with AB were HPAF-II, which presented a lower expression of *SPDEF* than CFPAC1 or BxPC3 and a similar expression to PATU8988S. We next explored *SPDEF* expression and mucus production in the same PDA cell lines and the slow-progressing human organoid line hF27 following orthotopic transplantation into NOD scid gamma mice (NSG) (Figure 3G, H). Notably, we found that PDA cells dramatically increased mucus production and fell into two groups: high SPDEF-expressing lines (hF27, HPAF-II and CFPAC1) and low SPDEF-expressing lines (BxPC3, PATU8988S, YAPC and AsPC1) (Figure 3G). High SPDEF-expressing lines hF27 and HPAF-II formed well-differentiated tumors with glandular lesions constituted by mucin-secreting columnar epithelial cells (Figure 3H). Tumors derived from the high SPDEF-expressing line CFPAC1 presented epithelial features but limited mucinous glands. CFPAC1 cells are unique as they were derived from a PDA patient with cystic fibrosis that carried a mutation resulting in the absence of a phenylalanine at position 508 in the cystic fibrosis transmembrane conductance regulator (delF508CFTR) (81). This mutation results in malformed CFTR protein production that leads to malfunction of a chloride channel and abnormalities in mucus secretion; thus, potentially explaining the different histology of CFPAC1 tumors from the other high SPDEF-expressing lines hF27 and HPAF-II.

On the other hand, low SPDEF-expressing lines formed highly cellular tumors with low mucus-secreting activity with the exception of PATU8988S, which gave rise to tumors with glandular architecture.

In conclusion, we found that high expression of the SPDEF program was associated with glandular differentiation and high secretory activity in PDA. Interestingly, these typical features of the classical subtype of PDA were retained by high SPDEF-expressing cell lines, but were only visible *in vivo*. On the other hand, low SPDEF-expressing cell lines formed poorly-differentiated tumors representative of the basal-like subtype.

### SPDEF can drive mucus production and promotes tumor growth of classical PDAs

To investigate if SPDEF supports the growth of human PDA, we deleted SPDEF in PDA cells using sgRNA pairs designed to eliminate the TSS and isolated single cell-derived clones. *SPDEF* inactivation was assessed at the mRNA level and the deletion of the TSS was confirmed by PCR on the genomic DNA (Figure S5A). Genetic manipulation of *SPDEF* did not have a clear effect on the proliferation of both high and low SPDEF-expressing PDA cells *in vitro* (Figure S5B, C). However, loss of *SPDEF* severely impaired the growth of high SPDEF-expressing cells hF27 and CFPAC1 *in vivo* following orthotopic transplantation and it extended the survival of orthotopically grafted HPAF-II tumor models (Figure 4A-C; S5D, E). On the other hand, deletion of *SPDEF* in low SPDEF-expressing PDA cell lines BxPC3, PATU8988S and YAPC did not affect tumor growth *in vivo* and, in AsPC1, it did not extend the survival of these orthotopically grafted tumor models (Figure 4D-G; S5F-H). Thus, SPDEF promoted tumor progression in high SPDEF-expressing tumors of classical differentiation but not in low SPDEF-expressing tumors of basal-like differentiation.

**Figure 4.**
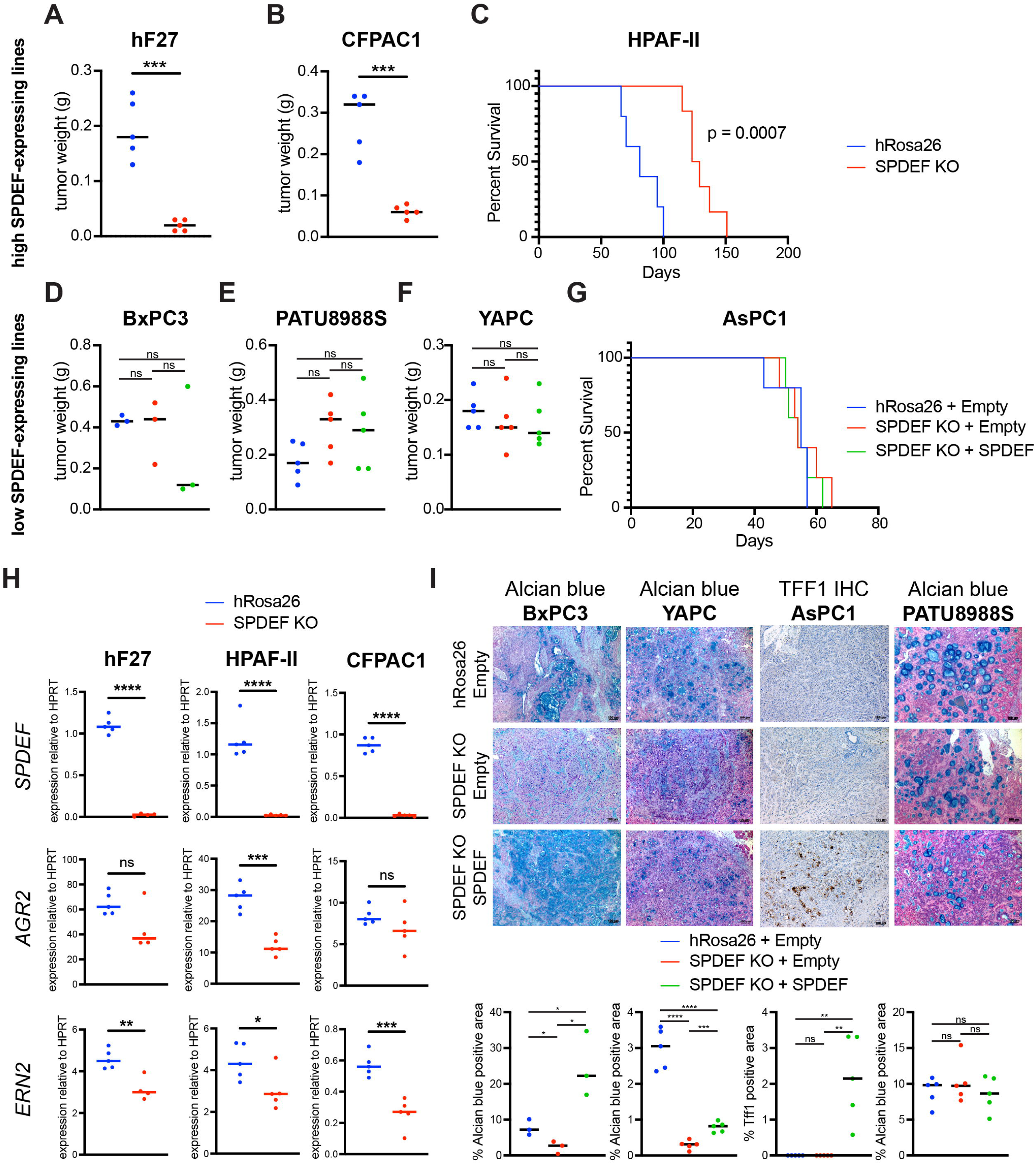
SPDEF can drive mucus production and promotes tumor growth of classical PDAs. **A, B**. Quantification of weight of tumors derived from hF27 (**A**) and CFPAC1 (**B**) orthotopically grafted models of *SPDEF* KO and hRosa26 clones in NSG mice. Results show mean of biological replicates. Unpaired Student’s t test. **C**. Kaplan–Meier survival curve of percent survival for HPAF-II orthotopically grafted models of *SPDEF* KO (n=5) and hRosa26 clones (n=5) in NSG mice. Log-rank (Mantel–Cox) test. **D-F**. Quantification of weight of tumors derived from BxPC3 (**D**), PATU8988S (**E**) and YAPC (**F**) orthotopically grafted models of hRosa26 and *SPDEF* KO clones with (SPDEF) or without (Empty) restoration of SPDEF expression in NSG mice. Results show mean of biological replicates. Unpaired Student’s t test. **G**. Kaplan–Meier survival curve of percent survival for AsPC1 orthotopically grafted models of hRosa26 (n=4) and *SPDEF* KO clones with (SPDEF) (n=5) or without (Empty) (n=5) restoration of SPDEF expression in NSG mice. **H**. *SPDEF, AGR2* and *ERN2* expression as determined by RT-qPCR in tumors derived from hF27, HPAF-II and CFPAC1 orthotopically grafted models of *SPDEF* KO and hRosa26 clones in NSG mice. Results show mean of biological replicates. Unpaired Student’s t test. **I**. Representative AB and nuclear red staining of BxPC3, YAPC, AsPC1 and PATU8988S orthotopically grafted models, in NSG mice, of hRosa26 and *SPDEF* KO clones with (SPDEF) or without (Empty) restoration of SPDEF expression. Scale bars, 100 μm. Bottom, average percentage of AB positive area. Each dot indicates a biological replicate. Unpaired Student’s t test.

Since the activation of the SPDEF program was associated with glandular epithelial differentiation and elevated secretory activity in PDA, we investigated whether deletion of SPDEF in high SPDEF-expressing tumors would affect their morphology and mucus-secreting activity. We found that loss of SPDEF did not alter the tumor histology: the tumors maintained their well-differentiated and glandular features (Figure S5I). In addition, to our surprise, AB staining did not show a reduction in mucus secretion in *SPDEF* knock-out tumors (Figure S5J). SPDEF is a known regulator of mucus production, so we wondered if these tumors epigenetically re-wired and re-expressed *SPDEF* or its target genes. However, we confirmed *SPDEF* inactivation by RT-qPCR on the tumor RNA and by RNA ISH on tumor sections (Figure 4H; S5K). Furthermore, we showed that SPDEF target genes *AGR2* and *ERN2* were down-regulated in *SPDEF* knock-out tumors, although inter-tumor variability across the cohorts was noted. Collectively, these observations indicated that SPDEF promoted the growth of well-differentiated glandular human tumors and may have done so by lowering the ER stress caused by mucus production through the regulation of the ER-resident proteins IRE1β and AGR2 among others, in accordance with our findings in murine epithelial^high^ pancreatic cancer cells. However, other pathways must compensate for SPDEF loss to regulate mucus production and secretion in human PDA.

To determine if SPDEF may promote mucus production in PDA, we performed gain-of-function experiments and over-expressed *SPDEF* in low SPDEF-expressing tumors. We analyzed mucus secretion by AB staining in orthotopically grafted BxPC3 and YAPC tumor models and found that deletion of *SPDEF* reduced mucus secretion in both models and *SPDEF* over-expression partially rescued mucus production in YAPC tumors, while it significantly induced mucus production in BxPC3 tumors (Figure 4I). AsPC1 control tumors were unique as they did not express *SPDEF* nor secrete mucus (see Figure 3G, H). In these tumors, SPDEF over-expression induced secretion of Tff1, a peptide found in mucus (Figure 4I). In AsPC1 tumors, mucus secretion was assessed by Tff1 IHC as necrotic areas were also AB positive, confounding the quantification. In conclusion, SPDEF regulated mucus production and secretion in low SPDEF-expressing tumors. Analysis of the tumor RNA found the expected expression trends: *SPDEF* and its targets *AGR2, ERN2, FOXA3 and MUC5AC* genes were down-regulated in *SPDEF* knock-out tumors and up-regulated in SPDEF over-expressing tumors, although inter-tumor variability in expression levels was observed (Figure S6). The only exception was the tumors derived from PATU8988S cells, which did not alter mucus production following *SPDEF* manipulation (Figure 4I). Different from the poorly-differentiated tumors derived from the transplantation of the other low SPDEF-expressing PDA cells, PATU8988S tumors were characterized by glandular differentiation and secretory activity.

Altogether, these findings supported the idea that SPDEF did regulate mucus production and secretion in human PDA. However, they also indicated that SPDEF was not the only regulator of epithelial and glandular identity in PDA cells, as SPDEF inactivation in glandular tumors did not alter the tumor differentiation and secretory activity.

## DISCUSSION

PDA is a heterogeneous disease (13,14,64). Here, we used a single-cell approach to determine the transcriptional changes underlying tumor progression in a well-established GEMM of pancreatic cancer. We identified multiple tumor subpopulations distinguished by their differentiation state and associated with different stages of the disease: from epithelial^high^ cancer cells to invasive mesenchymal and proliferative phenotypes. We found that the expression profiles associated with cell differentiation states in a mouse model were informative of human PDA subtypes. The current model for the evolution of human PDA subtypes indicates the classical phenotype as the default path of pathogenesis. The basal-like phenotype is selected for with disease progression in response to genetic aberrations, microenvironmental changes and epigenetic factors (13-15). Our data provided insight into the transcriptional changes associated with the acquisition of these phenotypes and revealed some of the transcription factors driving these phenotypes.

We discovered that Spdef was expressed by epithelial^high^ cancer cells in KPC tumors. Its expression was then reduced as tumor cells lost their epithelial features and became more aggressive. These findings were recapitulated in human PDA, where the expression of the SPDEF program was higher in classical rather than basal-like tumors and was associated with epithelial glandular differentiation. In addition, we showed that SPDEF promoted PDA tumors of epithelial/classical nature, which represent the majority of PDAs (7-9,11).

SPDEF was reported to exhibit both tumor-suppressive and oncogenic functions. Our current understanding of SPDEF’s role in mediating epithelial identity and supporting the fitness of epithelial pancreatic cancer cells could help to reconcile some of these controversial findings by considering the cell state. Indeed, SPDEF often displayed a tumor-promoting role in low-grade tumors with epithelial features and its expression was reduced during malignant differentiation. Re-expression of SPDEF in invasive tumors resulted in tumor-suppressive activity. For example, SPDEF induced mammary luminal epithelial lineage-specific gene expression and promoted the survival of luminal tumor cells (43). On the contrary, expression of SPDEF inhibited growth, motility and invasion of aggressive basal breast cancer cell lines (39,40). Similar to breast cancer cells, SPDEF promoted luminal epithelial differentiation in prostate cancer and deletion of SPDEF in luminal cells resulted in the induction of EMT-related proteins and increased migration while expression of SPDEF in invasive cells suppressed metastasis formation by inducing epithelial features (32-35,82). Altogether, these data indicated that SPDEF is an epithelial lineage specifier and not strictly an oncogene nor a tumor suppressor and its activity is dependent on the cell state.

Spdef is a well-defined regulator of the differentiation of secretory cells during development (23,28). In PDA, Spdef was expressed by mucinous neoplastic cells together with several other genes involved in mucus production, protein folding and glycosylation, some of which had been previously reported and confirmed by us to be regulated by Spdef. In addition, expression of the SPDEF program correlated with high secretory activity in human PDAs. We showed that SPDEF was sufficient but not necessary to regulate mucus production in PDAs. Indeed, expression of SPDEF in poorly-differentiated tumors induced mucus production. On the other hand, deletion of SPDEF in mucus-secreting tumors did not affect their secretory activity. Altogether, this indicated that SPDEF was one but not the only regulator of mucus production in PDA.

In addition to the regulation of mucus production, SPDEF supported the fitness of mucus-secreting tumors. This finding was supported by the discovery that SPDEF was not essential for the proliferation of PDA cells *in vitro*, but became a dependency *in vivo* for PDA of the classical subtype with a high secretory phenotype.

Highly secretory cells must adapt the activity of their ER to sustain the complex folding and glycosylation of secreted proteins (73). One of the strategies adopted by mucus-producing PDAs to deal with the stress caused by mucus production, is represented by the activation of the transcription factor MYRF, which maintains ER integrity and prevents ER stress by regulating genes involved in protein maturation and the unfolded protein response (UPR) (72). Here, we found that Spdef supported PDA growth by inducing the ER-resident disulfide isomerase Agr2 and ER stress sensor *Ern2*/Ire1β. Our data supported and extended the previous discovery that Agr2 is induced in response to ER stress and required for the initiation of pancreatic cancer (83). In addition, our study identified a role for *Ern2*/Ire1β in tumorigenesis. In Goblet cells and airway epithelium, *Ern2*/Ire1β, but not its most studied paralog and mediator of UPR *Ern1*/Ire1α, promoted efficient mucin production and folding in the ER (31,84-86). Our study indicated that the distinctive role of *Ern2*/Ire1β in preventing ER stress caused by the folding of mucins and other secreted proteins had a tumor-promoting role in PDA. Of note, *Ern1*/Ire1α inactivation is embryonic lethal, while *Ern2*/Ire1β mice are viable, indicating functional differences between the two paralogs (87-89). Our data suggest that the development of Ire1β-specific inhibitors could be beneficial for PDA of the classical subtype and potentially other mucinous tumors.

## ACKNOWLEDGEMENTS

We thank Dr. Taehoon Han for assistance with statistical analyses. This work was performed in collaboration with the Cold Spring Harbor Laboratory shared resources, which are supported by the National Institutes of Health (Cancer Center Support Grant 5P30CA045508: Bioinformatics, DNA Sequencing, Flow Cytometry, Animal, and Animal and Tissue Imaging Shared Resources). This work was performed with assistance from the US National Institutes of Health Grant S10OD028632-01.

The authors are supported by the Lustgarten Foundation, where D.A. Tuveson is a distinguished scholar and Director of the Lustgarten Foundation–designated Laboratory of Pancreatic Cancer Research. D.A. Tuveson is also supported by the Thompson Foundation, the Cold Spring Harbor Laboratory and Northwell Health Affiliation, the Northwell Health Tissue Donation Program, the Cold Spring Harbor Laboratory Association, and the National Institutes of Health (5P30CA45508, U01CA210240, R01CA229699, U01CA224013, 1R01CA188134, and 1R01CA190092). This work was also supported by a gift from the Simons Foundation (552716 to D.A. Tuveson). C. Tonelli was a fellow of the American-Italian Cancer Foundation. Y. Park is supported by the National Cancer Institute (R50CA211506).

## MATERIALS & METHODS

### Animals

*Kras*^*LSLG12D*^, *Trp53*^*LSLR172H*^ and *Pdx1-Cre* strains in C57Bl/6 background were interbred to obtain *Kras*^*LSLG12D/+*^; *Trp53*^*LSLR172H/+*^; *Pdx1-Cre* (KPC) mice (48,49). NOD scid gamma (NSG) mice were bred in house or purchased from The Jackson Laboratory; nu/nu mice were purchased from The Charles River Laboratory. All animal experiments were conducted in accordance with procedures approved by the IACUC at Cold Spring Harbor Laboratory.

### Pancreatic ductal organoid culture

Murine and human tumor organoids were established as described in (90). Cells were seeded in GFR Matrigel (BD). Once Matrigel was solidified, pancreatic organoid medium was added. Murine pancreatic organoid medium contains AdDMEM/F12, 10 mM HEPES (Invitrogen), Glutamax 1X (Invitrogen), penicillin/streptomycin 1x (Invitrogen), 500 nM A83-01 (Tocris), 50 ng/ml mEGF, 100 ng/ml mNoggin (Peprotech), 100 ng/ml hFGF10 (Peprotech), 0.01 mM hGastrin I (Sigma), 1.25 mM N-acetylcysteine, 10 mM Nicotinamide (Sigma), B27 supplement 1X (Invitrogen), R-spondin conditioned medium (10% final). Human pancreatic organoid medium contains AdDMEM/F12, 10 mM HEPES (Invitrogen), Glutamax 1X (Invitrogen), penicillin/streptomycin 1x (Invitrogen), 500 nM A83-01 (Tocris), 50 ng/ml hEGF, 100 ng/ml mNoggin (Peprotech), 100 μg/ml Primocin (InvivoGen), 100 ng/ml hFGF10 (Peprotech), 10 nM hGastrin I (Sigma), 1.25 mM N-acetylcysteine, 10 mM Nicotinamide (Sigma), B27 supplement 1X (Invitrogen), Wnt3a-conditioned medium (50% final), R-spondin conditioned medium (10% final).

### Cell lines culture

Human pancreatic cancer cells PATU8988S, HPAF-II, AsPC1, CFPAC1, YAPC and BxPC3 were cultured in RPMI supplemented with 5% fetal bovine serum (FBS). HEK293T cells were cultured in DMEM with 5% FBS. Penicillin/streptomycin were added to all cell cultures.

### Plasmids

Cas9-expressing pancreatic cancer cells were established by infection with LentiV_Cas9_puro vector (Addgene_108100). For knockout experiments, sgRNAs or sgRNA1-scaffold-bovineU6-sgRNA2 cassettes (synthesized as gene-blocks by IDT) were cloned into *Bsm*BI-digested LRG2.1_Neo (Addgene_125593). Sequences of sgRNAs are provided in Supplementary Table S3. The mouse *Spdef* cDNA and human *SPDEF* cDNA were synthetized from IDT and cloned into LentiV P2A Blast (Addgene_111887) using Gibson assembly (NEB).

### Virus production and transduction

Lentivirus was produced in HEK293T cells using helper plasmids (VSVG and psPAX2) with X-tremeGENE 9 DNA Transfection Reagent (Roche). Target cell lines were infected with the virus and 10 μg/ml polybrene. Organoids were dissociated into single cells by incubating them in TrypLE Express Enzyme (ThermoFisher) for 15 min while shaking and spin infected with the virus and 10 μg/ml polybrene (1700rpm for 45 min at room temperature). For organoid infection, the viral supernatant was concentrated 10 times with Lenti-X concentrator (Takara). Media were changed at 24 hours after infection and antibiotics (2 μg/ml puromycin, 1 mg/ml G418 or 10 μg/ml blasticidin) were added at 48 hours after infection.

### *In vitro* proliferation assay

Organoids were dissociated into single cells by incubating them in TrypLE Express Enzyme (ThermoFisher) for 15 min while shaking. Cells were counted and diluted to 10 cells/μl in a mixture of pancreatic organoid medium (90% final concentration) and GFR Matrigel (BD, 10% final concentration). 150 μl per well of this mixture (1500 cells per well) was plated in 96-well white plates (Nunc), whose wells had been previously coated with poly(2-hydroxyethyl methacrylate) (Sigma) to prevent cell adhesion to the bottom of the wells. Cell lines were trypsinized and counted. 500 cells/well were plated in 96-well white plates (Nunc) in RPMI supplemented with 5% FBS. Cell viability was measured every 24 hours, starting one day after plating, using the CellTiter-Glo assay (Promega) and SpectraMax I3 microplate reader (Molecular Devices). Five replicate wells per time point were measured.

### PCR-based analysis of TSS deletion

Genomic DNA from PDA cell lines or organoids was extracted with DNEasy Blood & Tissue Kit (Qiagen) following the protocol for cultured cells. Each PCR reaction was performed in a 20 μl mixture containing 1x AmpliTaq Gold 360 master mix (ThermoFisher), 0.5 μM each primer and 40 ng template DNA. Primer sequences are provided in Supplementary Table S3. The PCR cycling conditions were 95 °C for 5 minutes, followed by 30 cycles at 95 °C for 30 seconds, 60 °C for 30 seconds, and 72 °C for 30 seconds, with a final extension step at 72 °C for 5 minutes. PCR products were separated on a 2% agarose gel in 1x TAE buffer. Gel imaging was performed with a Syngene UV transilluminator.

### RNA Extraction

4-6 wells of organoids from a 24 well plate or 10^6^ PDA 2D cells or flash frozen tumor pieces were resuspended in 1 ml of TRIzol reagent. RNA was extracted using the TRIzol Plus RNA Purification Kit (ThermoFisher) per manufacturer’s instructions. RNA samples were treated on column with PureLink DNase (ThermoFisher).

### Expression analysis by RT-qPCR

Total RNA was isolated as described above. Complementary DNA (cDNA) was produced using the TaqMan reverse transcription reagents (ThermoFisher). 10 ng of cDNA were used for RT-qPCR reactions with TaqMan Universal Master Mix II, no UNG (ThermoFisher) and the following TaqMan probes: *Foxa3* Mm00484714_m1, *Spdef* Mm01306245_m1, *Agr2* Mm00507853_m1, *Ern2* Mm00469005_m1, *Hprt* Mm00446968_m1, *SPDEF* Hs00171942_m1, *AGR2* Hs00356521_m1, *ERN2* Hs01086607_m1, *FOXA3* Hs00270130_m1, *MUC5AC* Hs00873641_g1, *HPRT* Hs02800695_m1.

The expression of the *SPDEF* cDNA and the housekeeping gene GAPDH was analyzed with Power SYBR Green Master Mix (ThermoFisher) and the following primers: *For_SPDEF_cDNA* AGAAGACAGTTCTTGGGCGG, *Rev_SPDEF_cDNA* TGGGAGTCTATTACGGGGCA, *For_GAPDH* GGATTTGGTCGTATTGGG, *Rev_GAPDH* GGAAGATGGTGATGGGATT. *SPDEF* cDNA was codon optimized and could therefore be distinguished from endogenous *SPDEF*.

### RNA sequencing of Murine Organoids

RNA was extracted as described above. The quality of purified RNA samples was determined using a Bioanalyzer 2100 (Agilent) with an RNA 6000 Nano Kit. RNAs with RNA Integrity Number (RIN) values greater than 8.5 were used to generate sequencing libraries. Libraries were generated from 1 μg of total RNA using a TruSeq Stranded Total RNA Library Prep Human/Mouse/Rat (48 Samples) (Illumina) per manufacturer’s instructions. Libraries were quality checked using a Bioanalyzer 2100 (Agilent) with a High Sensitivity DNA Kit and quantified using PicoGreen (ThermoFisher). Equimolar amounts of libraries were pooled and subjected to paired-end, 75 bp sequencing at the Cold Spring Harbor DNA Sequencing Next Generation Shared Resource using an Illumina NextSeq 500 platform.

### RNA sequencing Data Analysis

RNA-seq reads quality was first quantified using FastQC version 0.11.8 (http://www.bioinformatics.babraham.ac.uk/projects/fastqc/). Reads were mapped to transcript annotation GENCODE M17 (91) using Spliced Transcripts Alignment to a Reference (STAR) version 2.5.2b (92). RSEM version 1.3.0 (93) was used to extract counts per gene. Differential gene expression analysis was performed using Bioconductor package DESeq2 version 1.24.0 (94). A pre-filtering step was performed to remove genes that have reads in less than two samples. At this step, all genes not classified as protein coding according to BioMart were discarded (95). Only genes with an adjusted p-value less than 0.05 were retained as significantly differentially expressed genes.

### *In vivo* transplantation assay

Organoids were cultured as described above and quickly harvested on ice in AdDMEM/F12 medium supplemented with HEPES 1x (Invitrogen), Glutamax 1x (Invitrogen), and penicillin/streptomycin (Invitrogen). Organoids were dissociated to single cells with TrypLE Express Enzyme (ThermoFisher). Cell lines were cultured as described above and trypsinized to single cells. Cells were resuspended in 50μl of GFR Matrigel (BD) diluted 1:1 with cold PBS. Mice were anesthetized using Isoflurane and subcutaneous administration of Ketoprofen (5 mg/kg). The surgery site was disinfected with Iodine solution and 70% ethanol. An incision was made in the upper left quadrant of the abdomen. 1 to 5*10^4^ murine tumor organoid cells were injected in the tail region of the pancreas of nu/nu mice; 2*10^5^ human tumor organoid cells or 10^4^ to 10^5^ human PDA cells were injected in the tail region of the pancreas of NSG mice. The incision at the peritoneal cavity was sutured with Coated Vicryl suture (Ethicon) and the skin was closed with wound clips (Reflex7, CellPoint Scientific Inc). Mice were euthanized and tumors were dissected out, weighted, photographed and processed for histology or RNA isolation. For the survival study, mice were taken when they reached a humane endpoint.

### Histology

All tissues were fixed with 10% neutral buffered formalin overnight. Tissues were processed with standard tissue processing protocol, embedded in paraffin and 6 μm sections were cut and mounted on slides. Formalin fixed paraffin-embedded (FFPE) tissue sections were stained with hematoxylin and eosin, Alcian blue (AB) and nuclear red or used for immunohistochemical staining. AB stain analysis was performed using ImageJ software for percentage area on a digital image of the tumor section acquired with Aperio scanner (Leica Biosystems) (96). Human pancreas tissue microarray was purchased from Biomax and used for AGR2 IHC (BIC14011a). The human PDA resection specimens used for ISH of *SPDEF* and *ERN2* were kindly provided by the Sidney Kimmel Comprehensive Cancer Center at Johns Hopkins University. All tissue donations and experiments were reviewed and approved by the Institutional Review Board of Cold Spring Harbor Laboratory and the clinical institutions involved. Written informed consent was obtained prior to acquisition of tissue from all patients. The studies were conducted in accordance with ethical guidelines (Declaration of Helsinki). Samples were confirmed to be tumor on pathologist assessment.

### Immunohistochemistry (IHC) and RNA in situ hybridization (RNA ISH)

FFPE sections were deparaffinized and rehydrated. For antigen retrieval, slides were incubated with 10 mM Citrate buffer (pH 6.0) in a pressure cooker for 6 minutes. To perform IHC, endogenous peroxidase activity was quenched in 3% H_2_O_2_ for 20 min. Tissues were blocked in 2.5% Normal Horse Serum blocking solution (Vector Laboratories) and subjected to staining with the following antibodies overnight at 4C: Fgfr2 (Abcam, ab58201 1:1000), Agr2 (CST, 13062 clone D9V2F 1:100), Gcnt3 (ThermoFisher, PA5-24455 1:100), Clca1 (Abcam, ab180851 1:100), Muc5ac (Abcam, ab3649 1:100), Gkn2 (LSBio, C294182 1:100), Gkn3 (LSBio, C294181 1:100), Tff1 (OriGene, TA322883 1:100), Tff2 (Proteintech, 13681-1-AP 1:100), Epcam (CST, 93790 clone E6V8Y 1:100). ImmPRESS Alkaline Phosphatase (AP) anti-Mouse IgG polymer and ImmPRESS Horseradish Peroxidase (HRP) anti-Rabbit IgG polymer, ImmPRESS HRP anti-Mouse IgG polymer (Vector Laboratories) were used as secondary antibodies for IHC. ImmPACT DAB peroxidase substrate and Vector Red Alkaline Phosphatase substrate (Vector Laboratories) were used as substrates. Hematoxylin (Vector Laboratories) was used as counterstain. Cover slides were mounted with Cytoseal 60.

RNA ISH was performed according to manufacturer’s instructions (RNAscope 2.5 HD Duplex Reagent Kit 322430 or RNAscope 2.5 HD Reagent Kit-RED 322350; Advanced Cell Diagnostics ACD) and probes specific for murine Spdef (544421-C2; ACD), murine Foxa3 (573031-C2; ACD), murine Ern2 (500431; ACD), murine Fgfr2 (443501; ACD), human SPDEF (543151, ACD) or human ERN2 (497231, ACD). Bright field images were obtained using a Zeiss Axio Imager.A2 microscope.

### Alcian blue and nuclear red staining on cell lines

Human pancreatic cell lines were seeded on 8-well tissue culture treated glass slide (Falcon). The day after, cells were fixed for 20 min with 4% paraformaldehyde at room temperature and then quenched with 3 consecutive incubations with 150 mM glycine for 5 min. Cells were washed with PBS and Alcian blue and nuclear red staining was performed with Alcian Blue (pH 2.5) Stain Kit (Vector) according to manufacturer’s instructions.

### Other bioinformatic and statistical analyses

GraphPad Prism 9 was used for graphical representation of data. Statistical analysis was performed using the tools within Prism 9 indicated in the figure legends. Asterisks denote P-value as follows: *, P ≤ 0.05, **, P ≤ 0.01, ***, P ≤ 0.001, ****, P ≤ 0.0001, ns, not significant. The SPDEF regulatory score was calculated by averaging the z-scores of *SPDEF, AGR2, ERN2, FOXA3* and *MUC5AC*. Z-scores were derived from log_2_(normalized counts) or log_2_(TPM values) as indicated in the figure legends. log_2_ values for unexpressed genes were set to 0. Heatmaps were generated using RStudio and Bioconductor package “ComplexHeatmap” (97,98). GO signatures were retrieved from GSEA MsigDB. Only genes expressed in at least two samples were analyzed.

Figures were prepared using Illustrator (Adobe).

### Tumor cells isolation

Primary KPC tumor tissues were carefully dissected avoiding adjacent normal pancreas or other tissue contamination. Tumors were minced and digested for 1 hour at 37C in digestion buffer (DMEM, 5% FBS, penicillin, streptomycin, 2.5 mg/ml Collagenase D (Sigma), 0.5 mg/ml Liberase DL (Sigma), 0.2 mg/ml DNase I (Sigma)) while shaking. Single-cell suspensions were obtained by filtering through 100 μm nylon cell strainers and subsequent hypotonic lysis of red blood cells using ACK lysis buffer (Gibco). KPC tumor cells were enriched with Tumor Cell Isolation Kit, mouse (Miltenyi Biotech) by magnetic cell sorting (MACS), according to manufacturer’s instructions.

### Single-cell RNA sequencing: library preparation and sequencing

Cancer cells from 8 KPC tumors were isolated as described above. Single cell suspensions with a viability higher than 80% were washed and resuspended in PBS supplemented with 0.5% bovine serum albumin (Sigma). Single-cell barcoded cDNA libraries were generated using the 10X Genomics Chromium Controller via the Single Cell 3’ Library Kit version 2 (120237, 10X Genomics). Libraries were sequenced on a NextSeq500 (Illumina) using the following custom run type: 26bp(R1)x56bp(R2)x8bp(i7) to a final depth of ∼60,000 mean reads per cell.

### Single-cell RNA sequencing: data processing

Reads were aligned to the mouse GRCm38 genome reference version 2.1.0 from 10X Genomics (cellranger-mm10-2.1.0) using Cellranger version 2.1.1 (10X Genomics) and default parameters.

Gene-cell matrices went through basic filtering to remove cells with fewer than 500 genes or more than 15% mitochondrial gene expressions. Cell normalization was performed using LogNormalize method in Seurat. Genes were scaled and centered after regression over percentage of mitochondrial and ribosomal expressions. Top 2000 highly variable genes (HVGs) were selected using Seurat version 2 method (99). In details, expression mean and dispersion were calculated for each gene across all cells and genes were grouped into bins based on their mean expression levels. Genes highly variant from other genes within the same bin were selected based on z-score normalized dispersion. HVGs were used to perform principal components analysis (PCA) and canonical correlation analysis (CCA) to align samples from multiple mice. After sample alignment, integrated analyses were performed including t-SNE dimensionality reduction and shared nearest neighbor (SNN) modularity optimization based cell clustering. Marker genes for each cluster were identified using ‘FindAllMarkers’ method available in Seurat package. Cell type identifications were carried out manually based on canonical cell marker genes. Contaminating non-tumor cell types were filtered from the data matrix. Only tumor cells were selected for further analysis. To further reveal cell heterogeneities within the tumor population, we obtained UMI count matrices for each mouse using cell barcodes identified from the above process and repeated the analysis using only tumor cells.

### Single-cell RNA-seq: pseudotime trajectory analysis

Construction of pseudotime trajectory was done mostly following Monocle 2 steps (50,100). The analysis started from UMI count matrix contains tumor cells. ‘negbinomial.size()’ was selected as expressionFamily, followed by size factor estimation, dispersion estimation, cell filtering and confounding factor regression, and cell clustering using unsupervised mode. ‘DDRTree’ was used for dimension reduction (101,102).

### Data availability

RNA-seq and scRNA-seq data generated in this study have been deposited in the Gene Expression Omnibus (GEO) database under the accession number GSE195502 and GSE195914, respectively.

Publicly available RNA-seq data reanalyzed for this study are available in the GEO database under the accession code GSE93326 (human specimen LCM RNA-seq), in the European Genome-phenome Archive (EGA) under study ID EGAS00001002543 (PanCuRx human specimen LCM RNA-seq), in the NCBI database of Genotypes and Phenotypes (dbGaP) under the accession number phs002045.v1.p1 (human organoids RNA-seq).

### Contact for reagents and resource sharing

Further information and requests for resources and reagents should be directed to the lead contact, Dr. David A. Tuveson.

## SUPPLEMENTARY FIGURE LEGENDS

**Figure S1.**
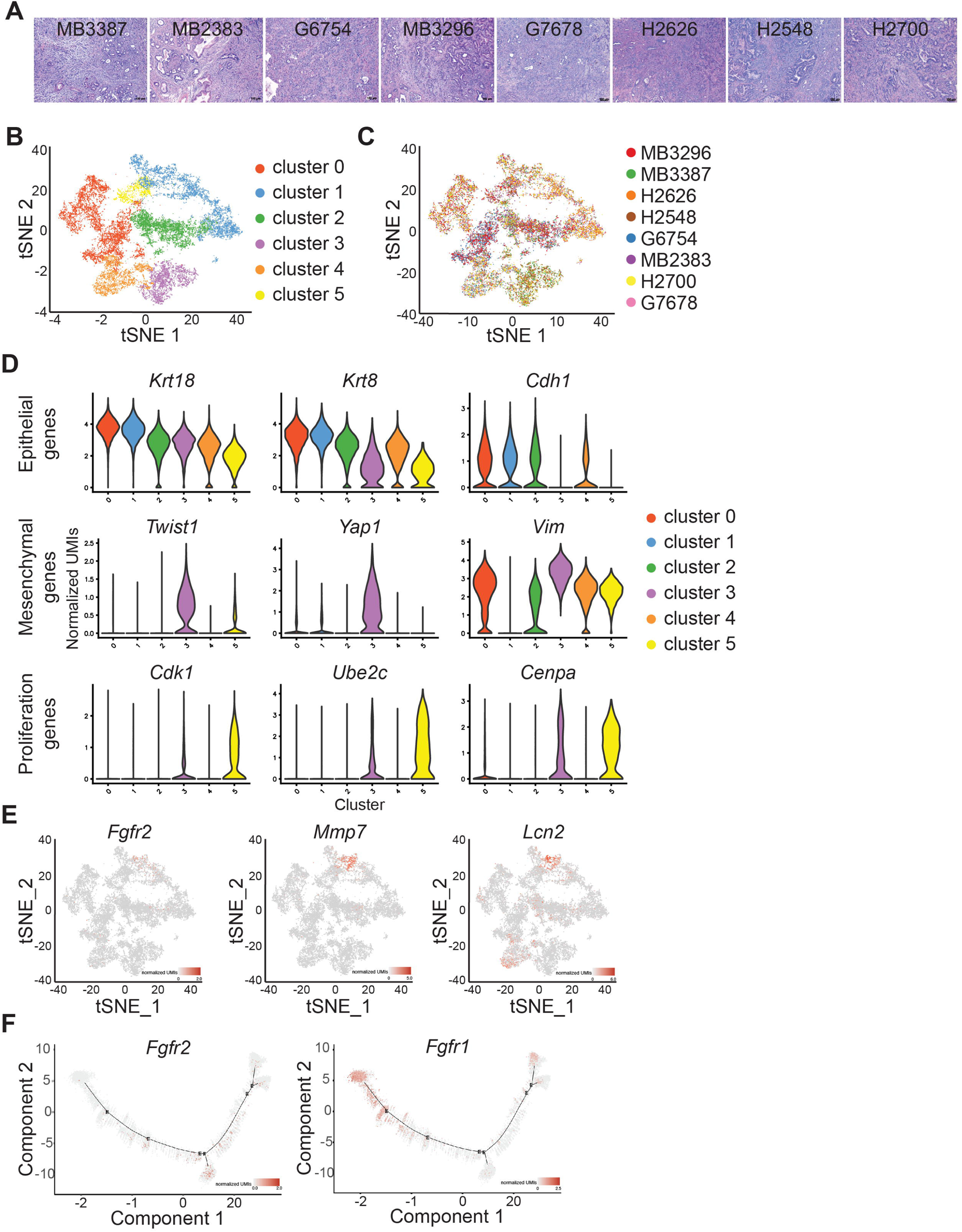
**A**. Hematoxylin and eosin (H&E) staining of the KPC tumors analyzed by scRNA-seq. Scale bars: 100 μm. **B**. tSNE plot of cancer cells, coloring by cluster. **C**. tSNE plot of cancer cells, coloring by sample. **D**. Violin plots showing the expression of epithelial, mesenchymal and proliferation genes in each cluster. **E**. Single-gene tSNE plots displaying the expression of *Fgfr2, Mmp7* and *Lcn2*. **F**. Single-gene pseudotime plots displaying the expression of *Fgfr2* and *Fgfr1*.

**Figure S2.**
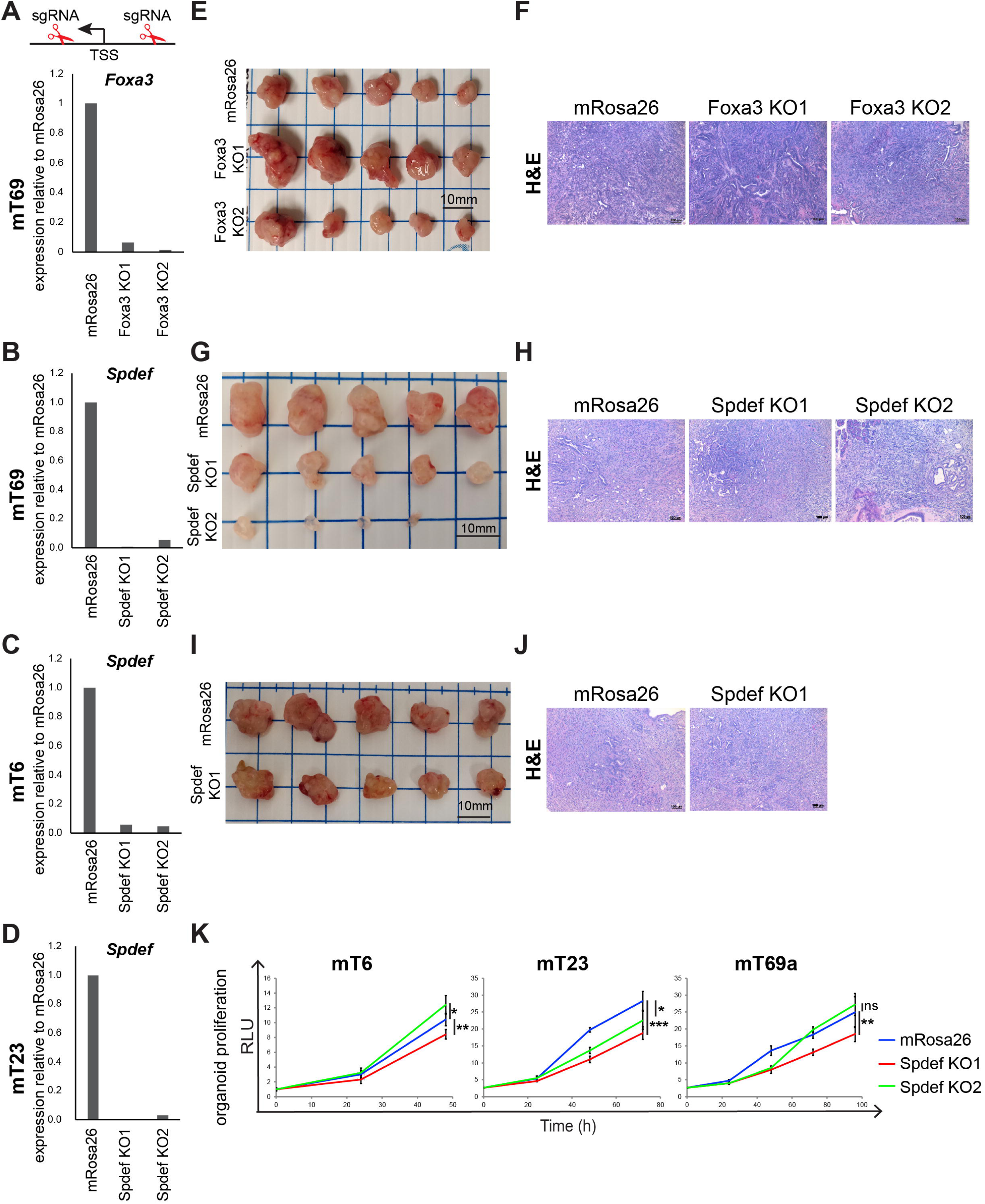
**A**. RT-qPCR analysis of *Foxa3* expression in *Foxa3* KO and mRosa26 clones in mT69a organoids. **B-D**. RT-qPCR analysis of *Spdef* expression in *Spdef* KO and mRosa26 clones in mT69a (**B**), mT6 (**C**) and mT23 (**D**) organoids. **E**. Images of tumors derived from mT69a OGO models of *Foxa3* KO and mRosa26 clones in nu/nu mice on day 26 after transplantation. **G, I**. Images of tumors derived from mT69a (**G**) and mT6 (**I**) OGO models of *Spdef* KO and mRosa26 clones in nu/nu mice on day 27 (**G**) and 30 (**I**) after transplantation. **F, H, J**. Representative H&E stain of tumors from E, G, I. Scale bars, 100 μm. **K**. Proliferation of *Spdef* KO and mRosa26 clones of mT6, mT23 and mT69a organoids measured with CellTiter-Glo luminescent cell viability assay. Unpaired Student’s t test.

**Figure S3.**
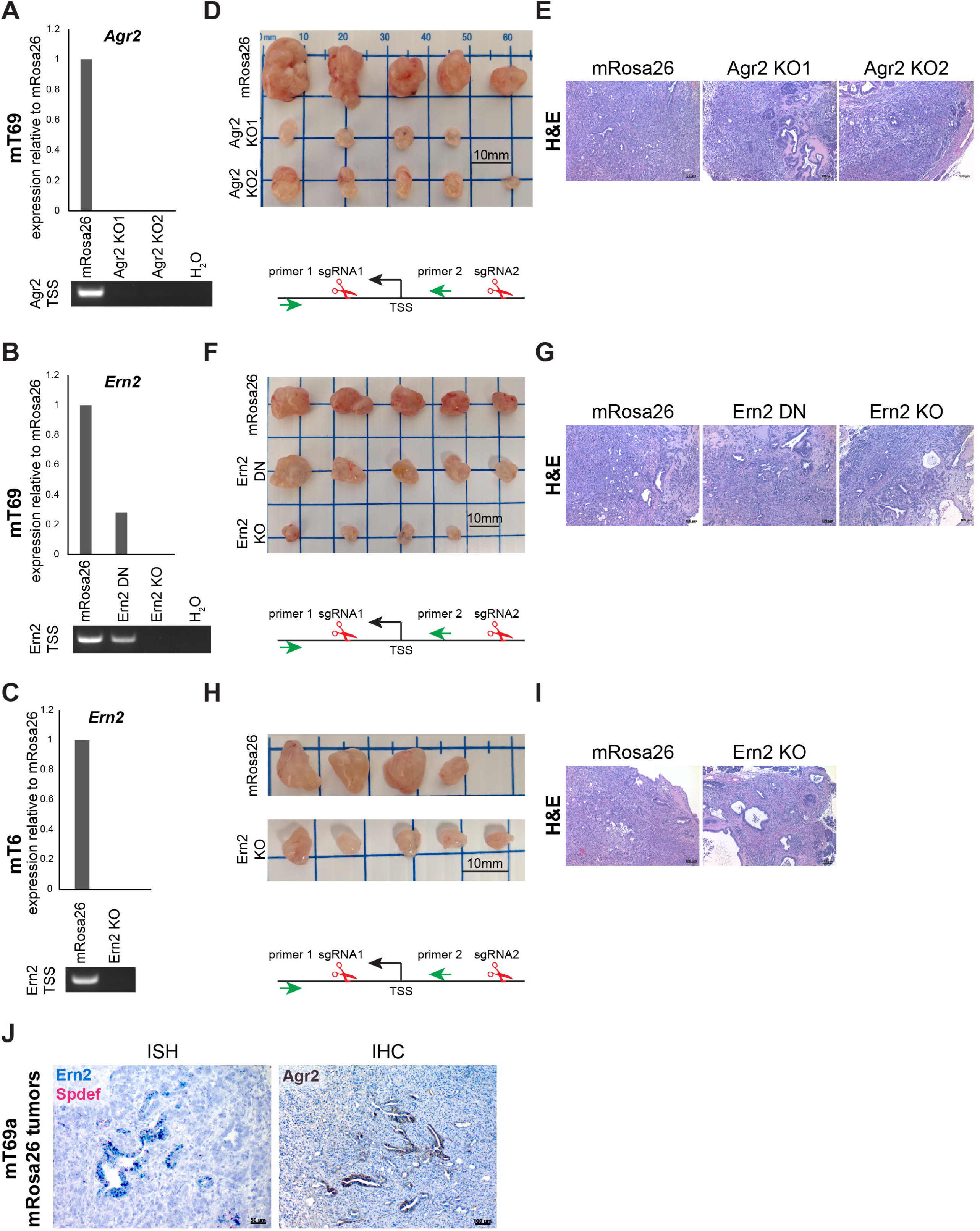
**A**. RT-qPCR analysis of *Agr2* expression and PCR-based analysis of TSS deletion in *Agr2* KO and mRosa26 clones in mT69a organoids. **B, C**. RT-qPCR analysis of *Ern2* expression and PCR-based analysis of TSS deletion in *Ern2* KO or partial inactivation DN and mRosa26 clones in mT69a (**B**), mT6 (**C**) organoids. **D**. Images of tumors derived from mT69a OGO models of *Agr2* KO and mRosa26 clones in nu/nu mice on day 26 after transplantation. **F, H**. Images of tumors derived from mT69a (**F**) and mT6 (**H**) OGO models of *Ern2* KO or partial inactivation DN and mRosa26 clones in nu/nu mice on day 26 (**F**) and 20 (**H**) after transplantation. **E, G, I**. Representative H&E stain of tumors from D, F, H. Scale bars, 100 μm. **J**. Representative RNA ISH of *Spdef* (red staining) and *Ern2* (blue staining) and immunohistochemical staining for Agr2 in tumor sections derived from mT69a OGO models of mRosa26 clones.

**Figure S4.**
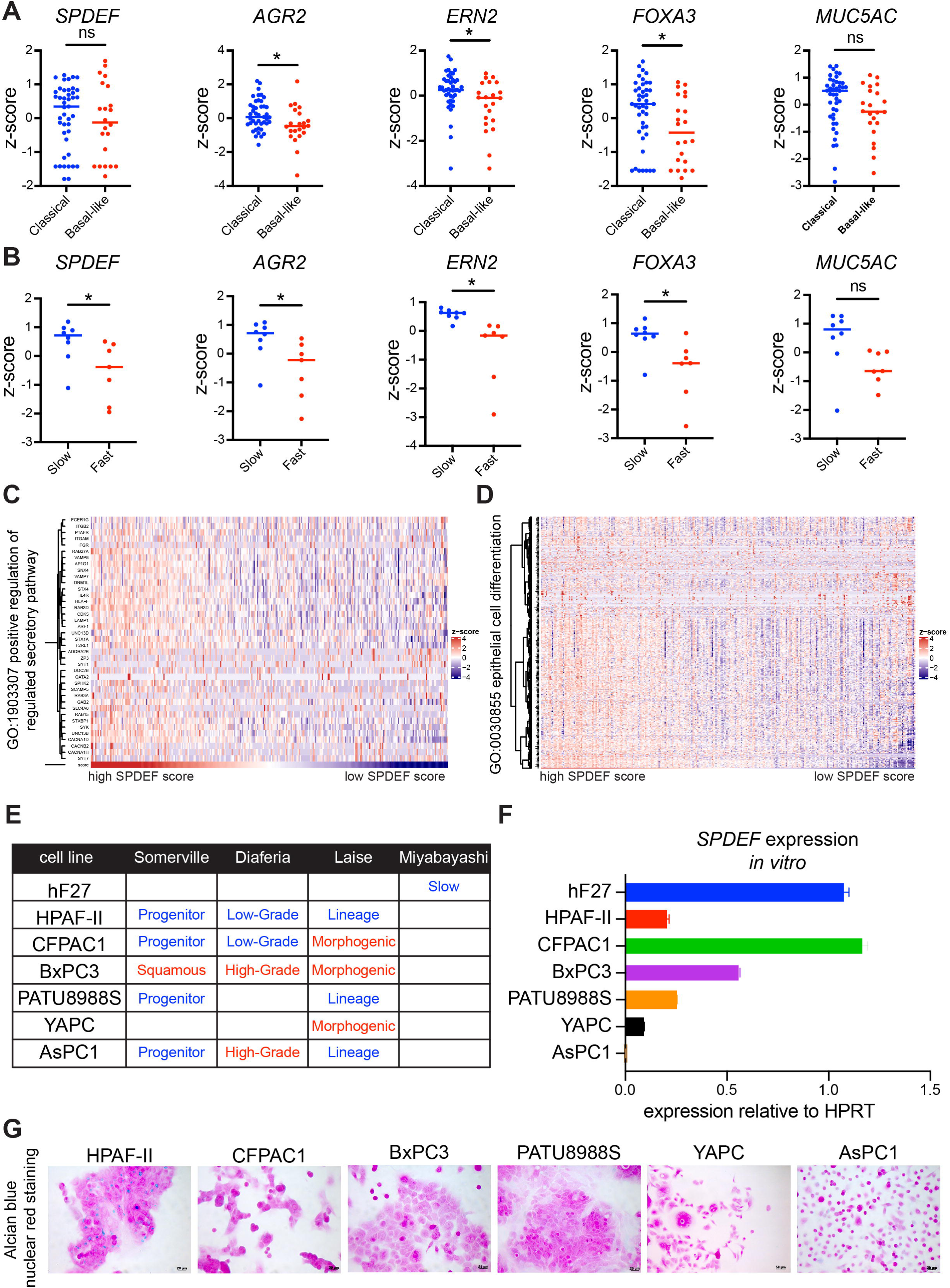
**A, B**. Z-score of log_2_(normalized counts) of *SPDEF, AGR2, ERN2, FOXA3* and *MUC5AC* in classical and basal-like PDA samples from (77) (**A**) and in Slow and Fast-progressing human organoid lines from (78) (**B**). **C, D**. Heatmap showing unbiased hierarchical clustering of RNA-seq data from the PanCuRx Translational Research Initiative dataset of patients with locally advanced PDA (79) for genes of GO signatures ‘positive regulation of regulated secretory pathway’ (**C**) and ‘epithelial cell differentiation’ (**D**). Samples are ranked based on their SPDEF score. The color scheme of the heatmap represents z-score of log_2_(TPM values). **E**. Classification of PDA cell lines based on their gene expression profiles in the indicated studies. **F**. RT-qPCR analysis of *SPDEF* expression in hF27, HPAF-II, CFPAC1, BxPC3, PATU8988S, YAPC, AsPC1 cell lines. Results show mean ± SD of two biological replicates. **G**. Representative AB and nuclear red staining of HPAF-II, CFPAC1, BxPC3, PATU8988S, YAPC, AsPC1 cell lines.

**Figure S5.**
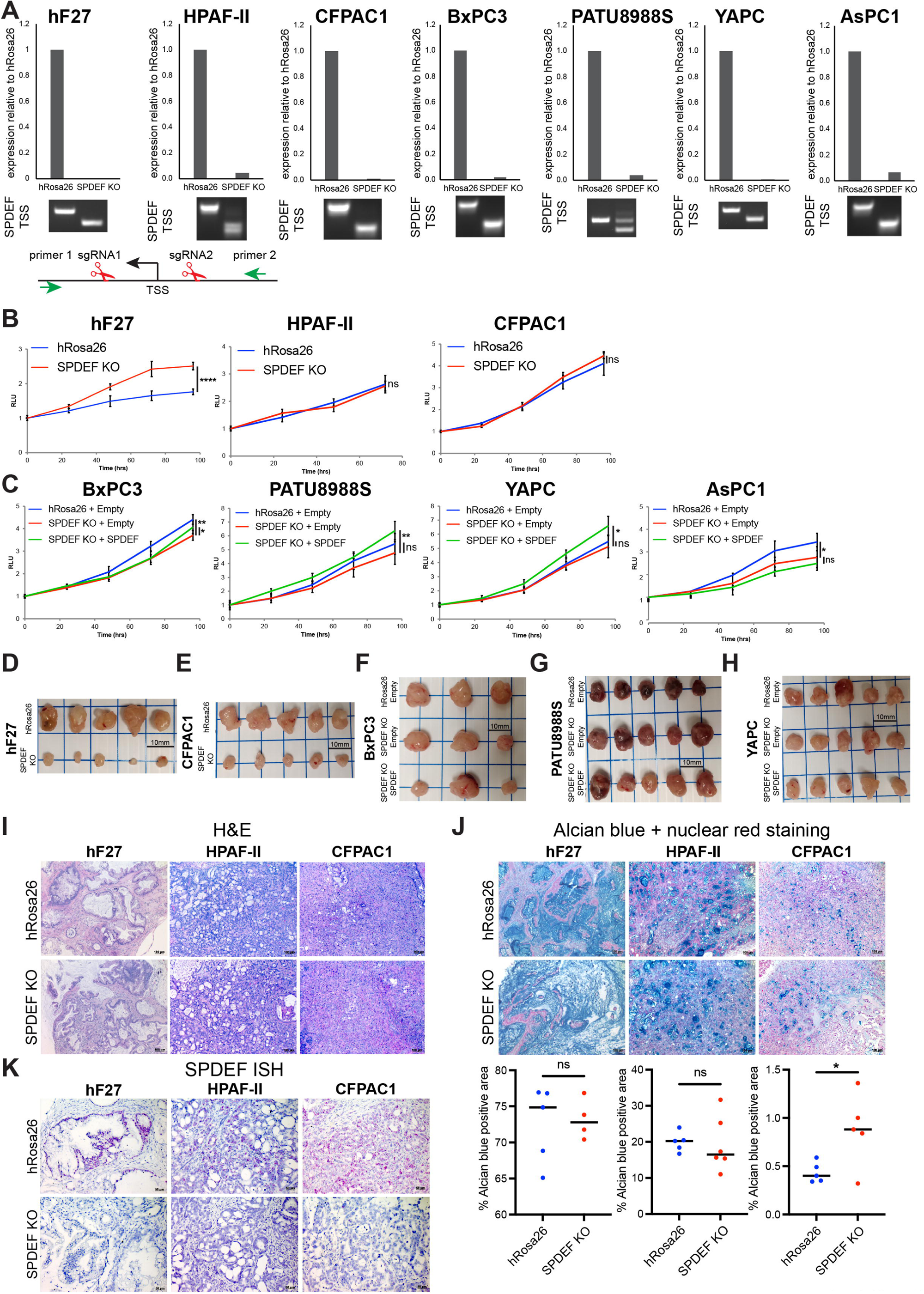
**A**. RT-qPCR analysis of *SPDEF* expression and PCR-based analysis of TSS deletion in *SPDEF* KO and hRosa26 clones in hF27 organoids and HPAF-II, CFPAC1, BxPC3, PATU8988S, YAPC, AsPC1 cell lines. **B**. Proliferation of hRosa26 and *SPDEF* KO clones in hF27 organoids and HPAF-II, CFPAC1 cell lines measured with CellTiter-Glo luminescent cell viability assay. Unpaired Student’s t test. **C**. Proliferation of hRosa26 and *SPDEF* KO clones with (SPDEF) or without (Empty) restoration of SPDEF expression in BxPC3, PATU8988S, YAPC, AsPC1 cell lines measured with CellTiter-Glo luminescent cell viability assay. Unpaired Student’s t test. **D, E**. Images of tumors derived from hF27 (**D**) and CFPAC1 (**E**) orthotopically grafted models of *SPDEF* KO and hRosa26 clones in NSG mice on day 217 (**D**) and 33 (**E**) after transplantation. **F-H**. Images of tumors derived from BxPC3 (**F**), PATU8988S (**G**) and YAPC (**H**) orthotopically grafted models of hRosa26 and *SPDEF* KO clones with (SPDEF) or without (Empty) restoration of SPDEF expression in NSG mice on day 55 (**F**), 43 (**G**) and 23 (**H**) after transplantation. **I**. Representative H&E stain of the tumors from Figure 4A-C. Scale bars, 100 μm. **J**. Representative AB and nuclear red staining of the tumors from Figure 4A-C. Scale bars, 100 μm. Bottom, average percentage of AB positive area. Each dot indicates a biological replicate. Unpaired Student’s t test. **K**. Representative RNA ISH of *SPDEF* in tumor sections derived from hF27, HPAF-II and CFPAC1 orthotopically grafted models of *SPDEF* KO and hRosa26 clones from Figure 4A-C. Scale bars, 50 μm.

**Figure S6.**
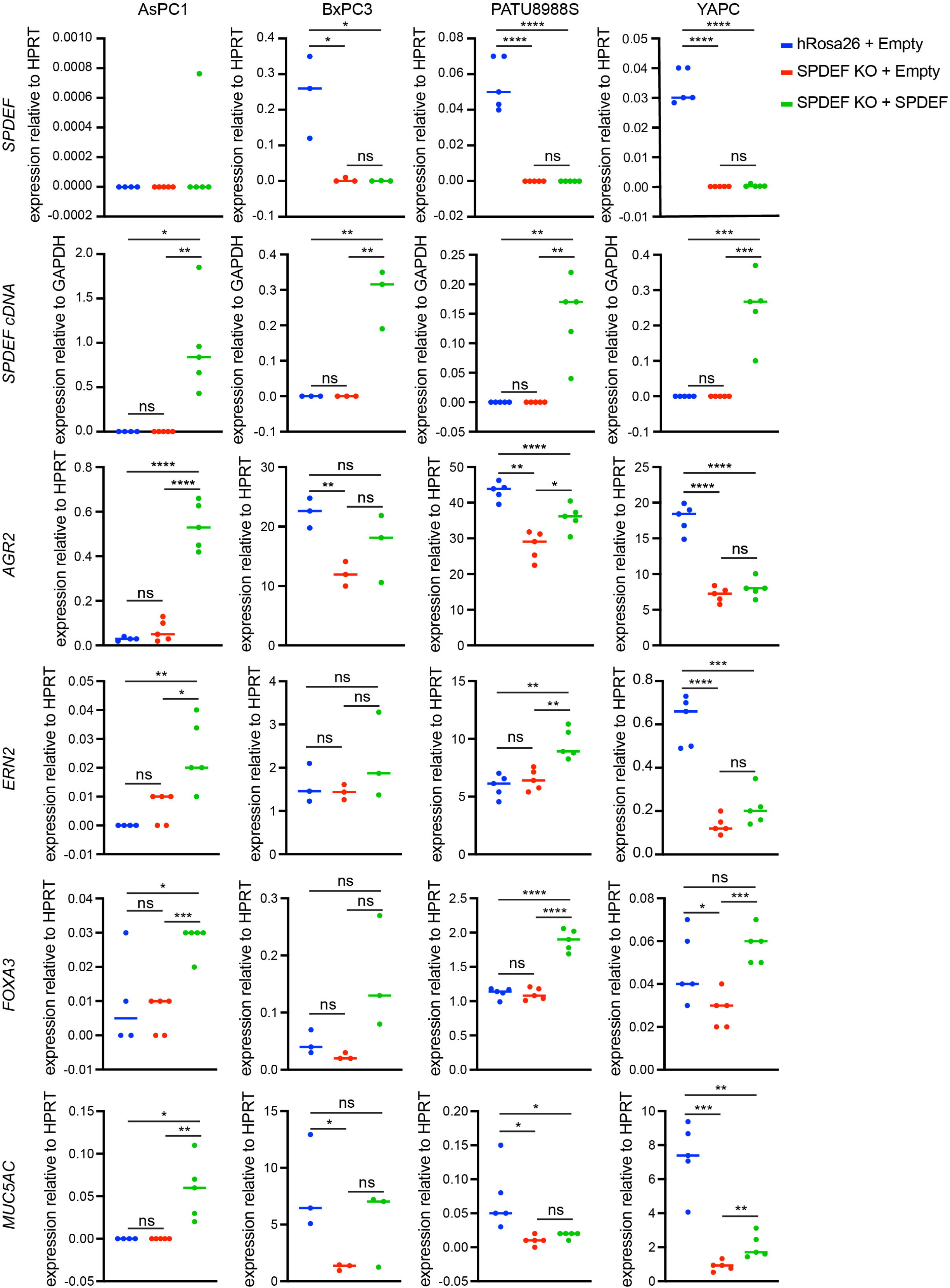
*SPDEF, SPDEF* cDNA, *AGR2, ERN2, FOXA3* and *MUC5AC* expression as determined by RT-qPCR in tumors derived from AsPC1, BxPC3, PATU8988S and YAPC orthotopically grafted models of hRosa26 and *SPDEF* KO clones with (SPDEF) or without (Empty) restoration of SPDEF expression in NSG mice. Results show mean of biological replicates. Unpaired Student’s t test.

